# FB5P-seq: FACS-based 5-prime end single-cell RNAseq for integrative analysis of transcriptome and antigen receptor repertoire in B and T cells

**DOI:** 10.1101/795575

**Authors:** Noudjoud Attaf-Bouabdallah, Iñaki Cervera-Marzal, Chuang Dong, Laurine Gil, Amédée Renand, Lionel Spinelli, Pierre Milpied

## Abstract

Single-cell RNA sequencing (scRNA-seq) allows the identification, characterization, and quantification of cell types in a tissue. When focused on B and T cells of the adaptive immune system, scRNA-seq carries the potential to track the clonal lineage of each analyzed cell through the unique rearranged sequence of its antigen receptor (BCR or TCR, respectively), and link it to the functional state inferred from transcriptome analysis. Here we introduce FB5P-seq, a FACS-based 5’-end scRNA-seq method for cost-effective integrative analysis of transcriptome and paired BCR or TCR repertoire in phenotypically defined B and T cell subsets. We describe in details the experimental workflow and provide a robust bioinformatics pipeline for computing gene count matrices and reconstructing repertoire sequences from FB5P-seq data. We further present two applications of FB5P-seq for the analysis of human tonsil B cell subsets and peripheral blood antigen-specific CD4 T cells. We believe our novel integrative scRNA-seq method will be a valuable option to study rare adaptive immune cell subsets in immunology research.

## Introduction

Technologies to reliably amplify and sequence the mRNA content of single-cells have progressed dramatically. By quantitatively measuring the expression levels of thousands of genes per cell, single-cell RNA sequencing (scRNA-seq) enables unbiased classification of cell types, and fine characterization of functional cell states^1,2^. Bioinformatics analyses of scRNA-seq datasets allow the identification of new cell types and functional subsets^3^, and reconstruct cellular differentiation or activation dynamics *a posteriori* from snapshot data^4–7^.

All scRNA-seq protocols are based on four common steps: (i) single-cell isolation, (ii) reverse transcription (RT) of mRNA, (iii) amplification of cDNA, and (iv) preparation of next-generation sequencing (NGS) libraries. Single-cell isolation can be performed through FACS or nanodroplet encapsulation. FACS has the advantage of allowing the user to record the precise cell surface phenotype of each sorted cell (index sorting) and link it to its deeply sequenced transcriptome (> 2,000 genes/cell), but with a limited throughput of a few hundred cells per sample^8,9^. Nanodroplet encapsulation, as performed in the commercial system proposed by 10x Genomics^10^, enables easy parallel processing of thousands of single-cells, albeit at the cost of reduced sensitivity (around 1,000 genes/cell in peripheral blood lymphocytes). Because, every scRNA-seq protocol have their own strengths and limitations^11,12^, method choice should be driven by the biological issue at hand but will also be constrained by the desired depth (number of genes detected per cell), throughput (number of cells analyzed per sample) and budget.

In the adaptive immune system, complex gene rearrangements generate diverse B cell receptor (BCR) and T cell receptor (TCR) repertoires enabling the recognition of an infinite range of antigens by B and T cells, respectively. When stimulated by their cognate antigen, each B or T cell clone can differentiate into multiple effector cell types that differ transcriptionally and functionally^13,14^. In the process of differentiation, the TCR sequence of mature T cells remains unchanged, while the BCR sequence of B cells may be altered in affinity maturation events such as class switch recombination and somatic hypermutation^15^. The progeny of a single T or B cell can thus be accurately identified through identical (for TCR) or very similar (for BCR) V-J junctional sequences in their TCR or BCR chain genes, respectively. Integrating single-cell immunoglobulin heavy chain (IGH) sequencing with low-throughput gene expression analysis by single-cell qPCR already revealed important features of memory B cell diversification^16^ and B cell lymphoma evolution^17^. Methods which enable parallel analysis of repertoire sequence and whole transcriptome gene expression in single B or T cells are required to deeply investigate multiple aspects of lymphocyte biology and malignancy.

Experimental and computational approaches to infer TCR and BCR sequence from scRNA-seq datasets of T and B cells have been developed, relying either on data produced by plate-based full-length sequencing (Smart-seq2)^18–22^, or droplet-based 5’-end sequencing (10x Genomics)^23^. The former allows for a deep analysis of phenotypically defined FACS-sorted cells, but is costly, labor intensive, and does not support unique molecular identifiers (UMIs). The latter incorporates UMIs, is relatively cheap and easy to perform on thousands of cells, but does not allow the precise selection of phenotypically defined cells and requires the production and sequencing of additional libraries for BCR or TCR sequencing.

Here we present FB5P-seq, a novel protocol for 5’-end scRNA-seq analysis of FACS-sorted cells, which incorporates UMI for accurate molecular counting and allows direct efficient recovery of paired BCR and TCR repertoire sequences when applied to B and T cells. We report the good sensitivity and accuracy of FB5P-seq when applied to human tonsil B cell subsets and antigen-specific peripheral blood T cells, highlighting the relevance and performance of our cost-effective and scalable technology.

## Results

### FB5P-seq experimental workflow

We based the design of the FB5P-seq experimental workflow on existing full-length^8^ and 5’-end^9,24^ scRNA-seq protocols. The main originalities in FB5P-seq were to perform cell-specific barcoding and incorporate 5 bp UMI during reverse transcription, and sequence the 5’-ends of amplified cDNAs from their 3’-end, and not from the transcription start site (**Figure 1A-B**). In FB5P-seq, single cells of interest are sorted in 96-well plates by FACS, routinely using a 10-color staining strategy to identify and enrich for specific subsets of B or T cells while recording all parameters through index sorting. Single-cells are collected in lysis buffer containing External RNA Controls Consortium (ERCC) spike-in mRNA (0.025 pg per well) and sorted plates are immediately frozen on dry ice and stored at −80°C until further processing. The amount of ERCC spike-in mRNA added to each well was optimized to yield around 5% of sequencing reads covering ERCC genes when studying lymphocytes which generally contain little mRNA. mRNA reverse transcription (RT), cDNA 5’-end barcoding and PCR amplification are performed with a template switching (TS) approach. Notably, our TSO design included a PCR handle (different from the one introduced at the 3’-end upon RT priming), an 8 bp well-specific barcode followed by a 5 bp UMI, a TATA spacer^25^, and three riboguanines. We empirically selected the 96 well-specific barcode sequences to avoid TSO concatemers in FB5P-seq libraries. After amplification, barcoded full-length cDNA from each well are pooled for purification and one-tube library preparation. For each plate, an Illumina sequencing library targeting the 5’-end of barcoded cDNA is prepared by a modified transposase-based method incorporating a plate-associated i7 barcode. The FB5P-seq library preparation protocol is cost-effective (260 € for library preparation of a 96-well plate), easily scalable and may be implemented on a pipetting robot.

**Figure 1.**
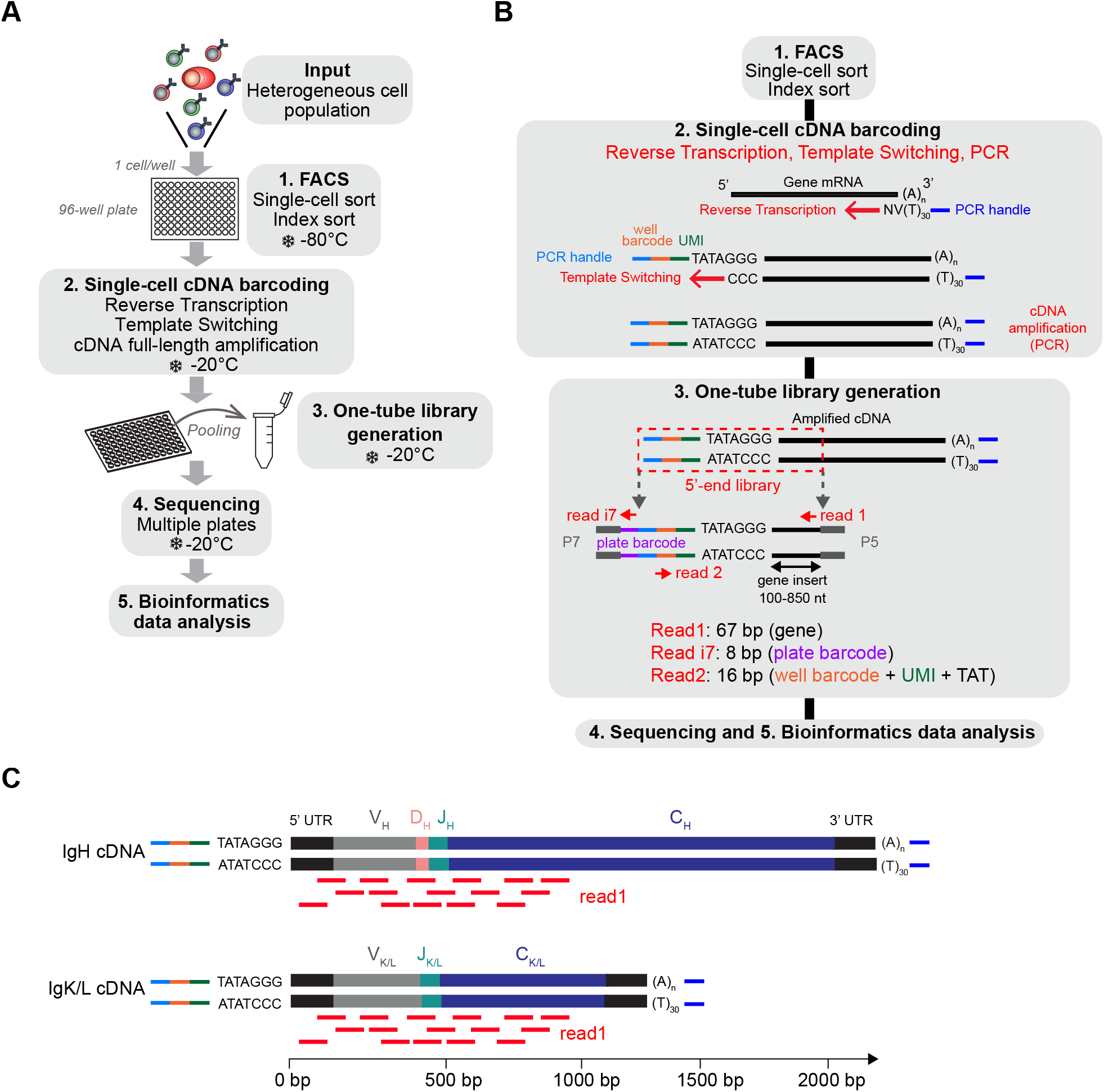
Overview of FB5P-seq experimental workflow. **(A)** Major experimental steps of the FB5P-seq workflow. **(B)** Schematic overview of molecular designs and reactions in the FB5P-seq workflow. **(C)** Schematic illustration of the mapping of Read1 sequences (in red) on IGH and IGK or IGL amplified cDNA, enabling the *in silico* reconstruction of paired variable BCR sequences.

FB5P-seq libraries are sequenced in paired-end single-index mode with Read1 covering the gene insert from its 3’-end, Read i7 assigning the plate barcode, and Read2 covering the well barcode and UMI. Because FB5P-seq libraries have a broad size distribution, with a gene insert of 100-850 bp, Read 1 sequences cover the 5’-end of transcripts approximately from 30 to 850 bases downstream of the transcription start site. Consequently, sequencing reads cover the whole variable and a significant portion of the constant region of the *IGH* and *IGK/L* expressed mRNAs (**Figure 1C**), enabling *in silico* assembly and reconstitution of BCR repertoire from scRNA-seq data. Because TCRα and TCRβ genes share a similar structure, FB5P-seq is equally suitable for reconstructing TCR repertoire from scRNA-seq data when T cells are analyzed.

### FB5P-seq bioinformatics workflow

The FB5P-seq data is processed to generate both a single-cell gene count matrix and single-cell BCR or TCR repertoire sequences when analyzing B cells or T cells, respectively. After extracting the well-specific barcode and UMI from Read2 sequences and filtering out low quality or unassigned reads, we use two separate pipelines for gene expression and repertoire analysis (**Figure 2**). The transcriptome analysis pipeline was derived from the Drop-seq pipeline^26^. Briefly, it consists of mapping all Read1 sequences to the reference genome, then quantifying, for each gene in each cell, the number of unique molecules through UMI sequences. After merging the data from all 96-well plates in the experiment, we filter the resulting gene-by-cell count matrices to exclude low quality cells, and normalize by total UMI content per cell (see Methods).

**Figure 2.**
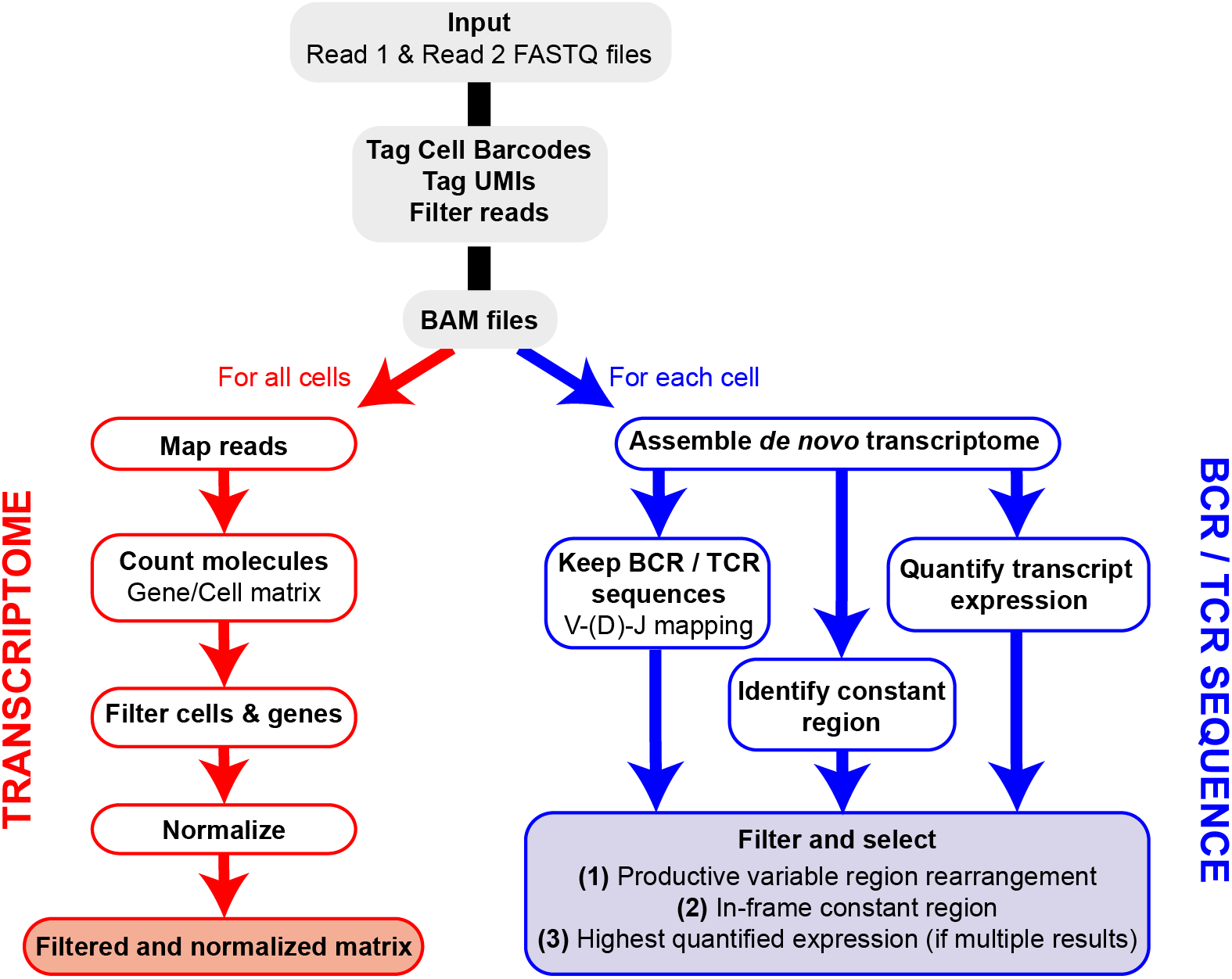
Overview of FB5P-seq bioinformatics workflow. Major steps of the bioinformatics pipeline starting from Read1 and Read2 FASTQ files for the generation of single-cell gene expression matrices and BCR or TCR repertoire sequences.

For the extraction of BCR or TCR repertoire sequences from FB5P-seq data, we have developed our own pipeline based on *de novo* single-cell transcriptome assembly and mapping of reconstituted long transcripts (contigs or isoforms) on public databases of variable immunoglobulin or TCR genes. We identify and select contigs corresponding to productive V(D)J rearrangements in-frame with an identified constant region gene. In cases where multiple isoforms are identified for a given chain (e.g. *IGH*) in a single cell, we assign the most highly expressed isoform and discard the other one(s). In early validation experiments, our pipeline was equally efficient and accurate as RT-PCR followed by Sanger sequencing for *IGH* variable region analysis (data not shown), with the major advantage of retrieving complete variable regions and large portions of constant regions of both *IGH* and *IGK/L*, or *TCRA* and *TCRB*, from the same scRNA-seq run.

### FB5P-seq quality metrics on human tonsil B cell subsets

To illustrate the performance of our scRNA-seq protocol, we obtained non-malignant tonsil cell suspensions from two adult human donors, referred to as Tonsil 1 and Tonsil 2. Based on surface marker staining, we excluded monocytes, T cells and naïve B cells, and sorted memory (Mem) B cells, germinal center (GC) B cells, and plasmablasts or plasma cells (PB/PCs) for FB5P-seq analysis (**Figure 3A**). We processed Tonsil 1 and Tonsil 2 samples in two separate experiments, generating libraries from 5 and 6 plates respectively. Libraries were sequenced at an average depth of approximately 500,000 reads per cell (**Table S1**). After bioinformatics quality controls (see Methods), we retained more than 90% of cells in the gene expression dataset (**Table S1**). We computed per cell accuracy (**Figure 3B**) and per experiment sensitivity (**Figure 3C**) based on ERCC spike-in detection levels and rates, respectively^11,12^. All cells showed high quantitative accuracy independently of their phenotype, with an overall mean correlation coefficient of 0.83 (**Figure 3B**). The molecular sensitivity ranged from 9.5 to 21.2 (**Figure 3C**), which compares favorably with other current scRNA-seq protocols^11,12^. We detected a mean of 987, 1712 and 1307 genes per cell in Mem B cells, GC B cells and PB/PCs, respectively (**Figure 3D**). GC and Mem B cells displayed higher total molecule counts (mean UMI counts of 192,765 and 145,356, respectively) than PB/PCs (mean UMI count of 67,861) (**Figure 3E**).

**Figure 3.**
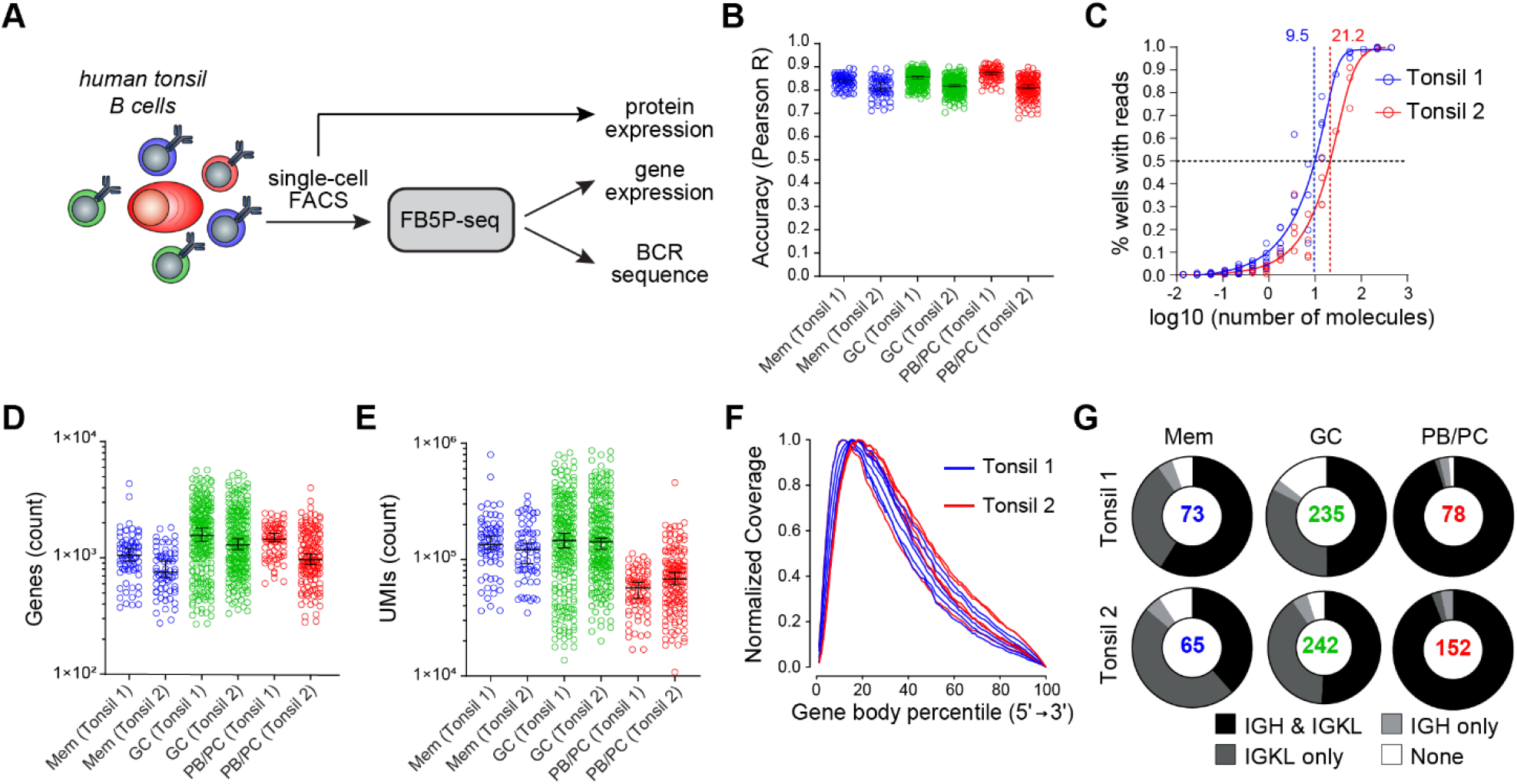
FB5P-seq quality metrics on human tonsil B cell subsets. **(A)** Experimental workflow for studying human tonsil B cell subsets with FB5P-seq. **(B)** Per cell quantitative accuracy of FB5P-seq computed based on ERCC spike-in mRNA detection (see Methods) for Memory B cells (Mem, n=73 Tonsil 1, n=65 Tonsil 2), GC B cells (GC, n=235 Tonsil 1, n=242 Tonsil 2) and PB/PC cells (n=78 Tonsil 1, n=152 Tonsil 2). Black line indicates the median with 95% confidence interval error bars. **(C)** Molecular sensitivity of FB5P-seq computed on ERCC spike-in mRNA detection rates (see Methods) in two distinct experiments. Dashed lines indicate the number of ERCC molecules required to reach a 50% detection probability. **(D-E)** Total number of unique genes **(D)** and molecules **(E)** detected in human tonsil Mem B cells (n=73 Tonsil 1, n=65 Tonsil 2), GC B cells (n=235 Tonsil 1, n=242 Tonsil 2) and PB/PCs (n=78 Tonsil 1, n=152 Tonsil 2). Black line indicates the median with 95% confidence interval error bars. **(F)** Gene body coverage analysis for Tonsil 1 (n=5) and Tonsil 2 (n=6) plate libraries. Each curve was computed from the BAM file corresponding to one library from a 96-well plate pool (see Methods). **(G)** Pie charts showing the relative proportion of cells with reconstructed productive IGH and IGK/L sequences (black), only IGK/L sequences (dark grey), only IGH sequences (light grey) or no BCR sequence (white) among Mem B cells, GC B cells and PB/PC cells from Tonsil 1 and Tonsil 2 samples. Total number of cells analyzed for each subset is indicated at the center of the pie chart.

As expected from the method design, FB5P-seq Read1 sequence coverage was biased towards the 5’-end of gene bodies, with a broad distribution robustly covering from the 3^rd^ to the 60^th^ percentile of gene body length on average (**Figure 3F**). In Tonsil 1 and Tonsil 2 B cell subsets, the BCR reconstruction pipeline retrieved at least one productive BCR chain for the majority of the cells (**Figure 3G**). Consistent with high expression of BCR gene transcripts for sustained antibody production, we obtained the paired *IGH* and *IGK/L* repertoire for the vast majority of PB/PCs. In Mem and GC B cells, we obtained paired *IGH* and *IGK/L* sequences on approximately 50% of the cells, and only the *IGK/L* sequence in most of the remaining cells. The superior recovery of *IGK/L* sequences was likely because the expression level of *IGK/L* was about 2-fold higher than *IGH* in our FB5P-seq data (data not shown).

Altogether, accuracy, sensitivity, gene coverage and BCR sequence recovery highlighted the high performance of the FB5P-seq method for integrative analysis of transcriptome and BCR repertoire in single B cells.

### FB5P-seq analysis of human tonsil B cell subsets

As a biological proof-of-concept, we further analyzed the Tonsil 1 and Tonsil 2 datasets. T-distributed stochastic neighbor embedding (t-SNE) analysis on the gene expression data discriminated three major cell clusters. Tonsil B cells clustered based on their sorting phenotype (Mem B cells, GC B cells or PB/PC) and did not cluster by sample origin (**Figure S1A**). Cell cycle status further separated the cycling (S and G2/M phase) from the non-cycling (G1) GC B cells (**Figure S1B**). The expression levels of surface protein markers recorded through index sorting were consistent with the gating strategy of Mem B cells (CD20^+^CD38^lo^CD10^-^CD27^+^), GC B cells (CD20^+^CD38^+^CD10^+^) and PB/PCs (CD38^hi^CD27^hi^) (**Figure 4A**, top panel). The expression of the corresponding mRNAs mirrored the protein expression (**Figure 4A**, bottom panel), but revealed numerous cells where the mRNA was undetected despite intermediate or high levels of the protein. Further, we detected the expression of known marker genes for Mem B cells (*CCR7, SELL, GPR183*) GC B cells (*AICDA, MKI67, CD81*) or PB/PC (*XBP1, PRDM1, IRF4*) in the corresponding clusters (**Figure 4B**), and identified the top marker genes for each cell subset (**Figure 4C**). Those analyses were consistent with previous single-cell qPCR analyses^17^ and bulk microarray analyses of human B cell subsets^27,28^.

**Figure 4.**
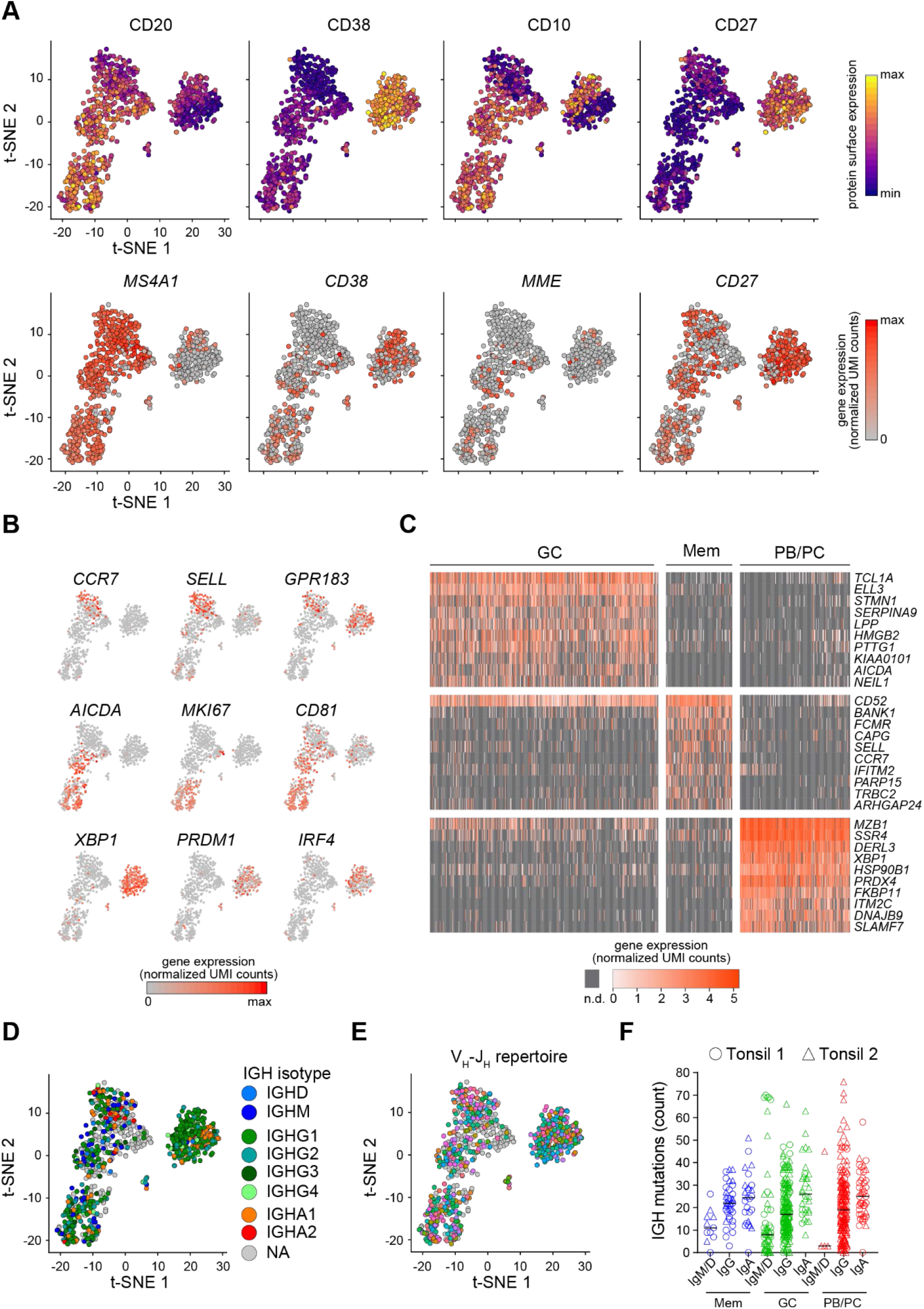
FB5P-seq analysis of human tonsil B cell subsets. **(A)** t-SNE map of single human tonsil B cell subsets computed on 4,000 variable genes excluding BCR genes. Cells are colored based on surface protein (*upper panel*) or corresponding gene (*lower panel*) expression of the indicated markers (n=845 cells). **(B)** Expression of indicated marker genes for Mem B cells (*upper panel*), GC B cells (*middle panel*) or PB/PCs (*bottom panel*) in single human tonsil B cells laid out in the t-SNE map. **(C)** Gene expression heatmap of GC B cells (n=477), Mem B cells (n=138) and PB/PCs (n=230) for the top 10 marker genes of each subset (n.d.: not detected). **(D-E)** t-SNE map of single human tonsil B cell subsets colored by IGH isotype (NA: not applicable i.e. no IGH reconstructed) **(D)** or V_H_-J_H_ repertoire (grey cells: no IGH reconstructed) **(E)**. **(F)** Scatter plots showing IGH mutation frequency in human Tonsil 1 (circles) and Tonsil 2 (triangles) B cells sorted by their IGH isotype and phenotype (Mem B cells: n=11 IgM/IgD+, n=37 IgG+ and n=26 IgA+; GC B cells: n=55 IgM/IgD+, n=174 IgG+ and n=32 IgA+; PB/PC: n=4 IgM/IgD+, n=179 IgG+ and n=42 IgA+ PB/PCs. Black line indicates the median.

Integrating the single-cell BCR repertoire data to the t-SNE embedding, we revealed that the *IGH* and *IGK/L* repertoire of tonsil B cell subsets was polyclonal (**Figure 4D-E** and **Figure S1C-D**). Interestingly, while the somatic mutation load was equivalent in Igκ and Igλ light chains from Mem B cells, GC B cells and PB/PCs (**Figure S1E**), the *IGH* mutation rate depended on isotype, with IgA^+^ cells expressing the most mutated BCR (**Figure 4F**) regardless of phenotype or sample origin. By contrast, IgM/IgD^+^ cells exhibited the lowest somatic mutation loads (**Figure 4F**).

Overall, those analyses confirmed that the FB5P-seq method is relevant for simultaneous protein, whole-transcriptome and BCR sequence analysis in human B cells.

### FB5P-seq analysis of human peripheral blood antigen-specific CD4 T cells

To test whether our protocol is also effective in T cells, we applied FB5P-seq to *Candida albicans*-specific human CD4 T cells sorted after a brief restimulation of fresh peripheral blood mononuclear cells with a pool of MP65 antigen-derived peptides^29^ (**Figure 5A** and Methods). *Candida albicans* is a common commensal in humans, known to generate antigen-specific circulating memory CD4 T cells with a TH17 profile^30^. Similar to the B cell dataset, the T cell dataset displayed high per cell accuracy (**Figure 5B**) and an average of 1890 detected genes per cell (**Figure 5C**). Gene expression analysis showed an efficient detection of T cell marker genes (*CD3E*), activation genes (*CD40LG, EGR2, NR4A1, IL2*), and TH17-specific genes (*CCL20, CSF2, IL22, IL23A, IL17A*) in those reactivated antigen-specific T cells (**Figure 5D**). We recovered at least one productive TCRα or TCRβ chain in 88% of cells, and paired TCRαβ repertoire in 61% of cells (**Figure 5E**). Moreover, CDR3β sequence analysis revealed some expanded TCRβ clonotypes likely related to MP65 antigen-specificity (**Figure 5F**). Principal Component Analysis (PCA) of the gene expression data and visualization of V_β_-J_β_ TCR rearrangements revealed no apparent segregation of antigen-specific T cells expressing different clonotypes (**Figure 5G**).

**Figure 5.**
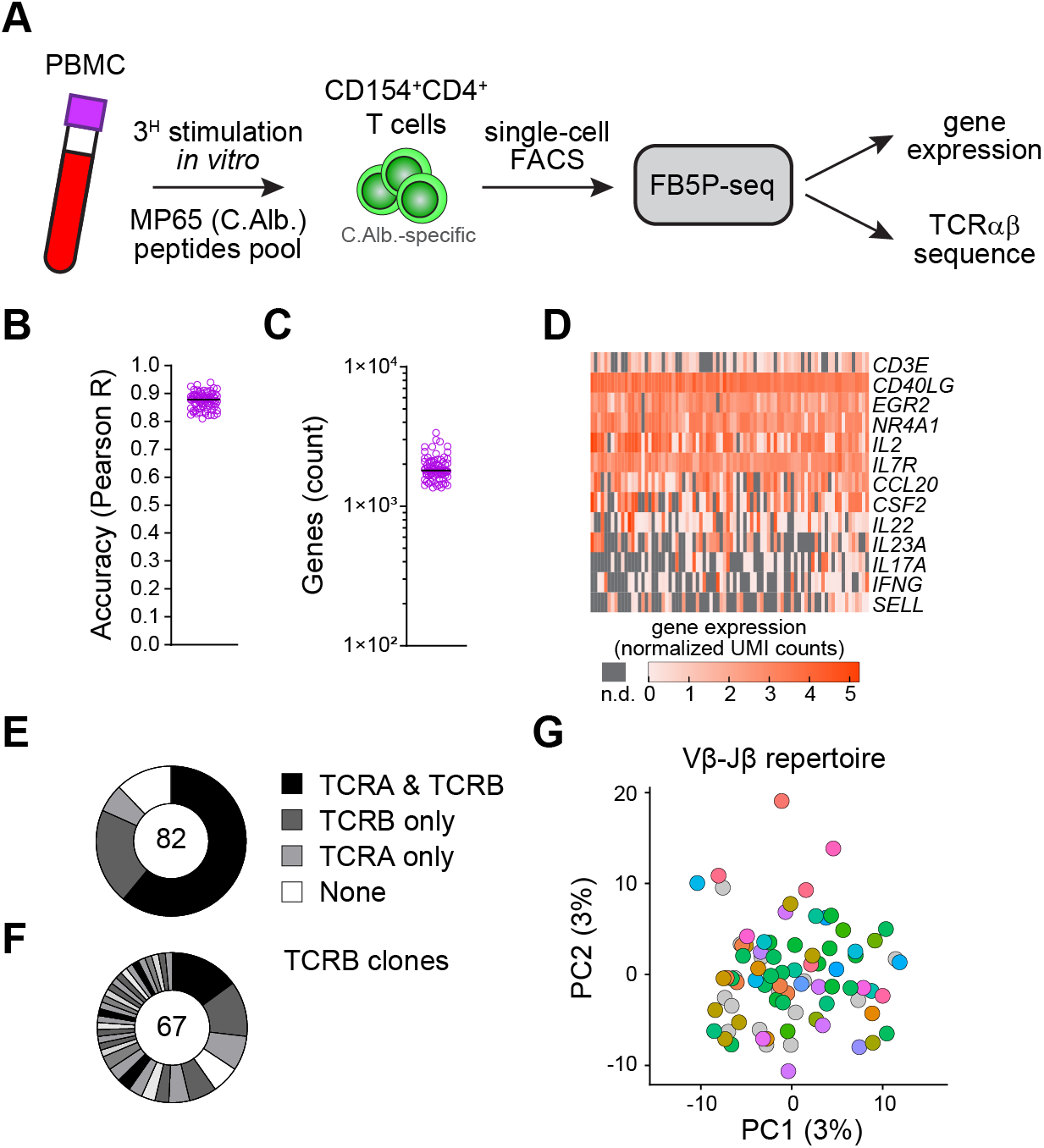
FB5P-seq analysis of human peripheral blood antigen-specific CD4 T cells. **(A)** Experimental workflow for studying human peripheral blood *Candida albicans*-specific CD4 T cells with FB5P-seq. **(B)** Per cell quantitative accuracy of FB5P-seq computed based on ERCC spike-in mRNA detection (see Methods) for *Candida albicans*-specific CD4 T cells (n=82). Black line indicates the median with 95% confidence interval error bars. **(C)** Total number of unique genes detected in *Candida albicans*-specific CD4 T cells (n=82). Black line indicates the median with 95% confidence interval error bars. **(D)** Gene expression heatmap of human peripheral blood *Candida albicans-specific* CD4 T cells for a selected panel of marker genes (n.d. : not detected). **(E)** Pie charts showing the relative proportion of cells with reconstructed productive TCRA and TCRB sequences (black), only TCRB sequences (dark grey), only TCRA sequences (light grey) or no TCR sequence (white) among *Candida albicans*-specific CD4 T cells (n=82). **(F)** Distribution of TCRB clones among *Candida albicans*-specific CD4 T cells (n=67). Black and grey sectors indicate the proportion of TCRB clones (clonotype expressed by ≥ 2 cells) within single-cells analyzed (white sector: unique clonotypes). **(G)** Projection of *Candida albicans*-specific CD4 T cells (n=82) on the first 2 PCs computed on 4,000 variable genes excluding TCR genes (PC1 : 3% of total variability, PC2 : 3% of total variability). Cells are colored based on V_β_-J_β_ repertoire (grey cells: no TCRB reconstructed).

Taken together, these data indicate that our method is also relevant for integrative scRNA-seq analysis of human T cells, especially for tracking rare antigen-specific cells *ex vivo*.

## Discussion

FB5P-seq is a novel 5’-end scRNA-seq workflow that allows accurate gene expression analysis of hundreds of FACS-sorted cells in parallel. When applied to B or T cells, the FB5P-seq data supports *in silico* reconstruction of paired full-length BCR or TCR variable repertoire on a per cell basis. As demonstrated by our studies of human tonsil B cells and peripheral blood antigen-specific T cells, FB5P-seq is particularly adapted for the integrative analysis of surface phenotype (through index sorting), gene expression, and antigen receptor repertoire of rare phenotypically defined B or T cells. Because FB5P-seq integrates three layers of barcodes (UMIs, cellular barcodes, plate barcodes), the workflow is more cost- and time-effective than Smart-Seq2^31^, which so far has been the method of choice for plate-based scRNA-seq analysis of gene expression and antigen receptor repertoire in B or T cells^18,19,21,22^. The molecular detection limit of FB5P-seq (10 to 20 molecules) was higher than what has been reported for Smart-seq2^11^ (7 molecules), suggesting that FB5P-seq may recover the expression of less genes per cell. In a recent benchmarking study of scRNA-seq methods, Smart-Seq2 detected the expression of approximately 2,500 genes per cell in human PBMCs at a sequencing depth of 1×10^6^ reads per cell^32^. The sensitivity of FB5P-seq was sufficient to recover in the order of 1,000 to 2,000 genes per cell on average in human lymphocytes (depending on cell type and cell cycle status) at a sequencing depth of approximately 5×10^5^ reads per cell.

One of our main constraints when developing FB5P-seq was to enable the *in silico* reconstruction of BCR or TCR repertoire sequences directly from the scRNA-seq reads. Because the variable regions of antigen receptor genes are encoded from the 5’-end of mRNAs, 3’-end scRNA-seq protocols are not suitable for parallel analysis of BCR or TCR repertoire, unless a separate amplification with cell-specific and gene-specific primers is performed on full-length cDNA before pooling single-cell contents^33^. Full-length scRNA-seq protocols with paired-end sequencing have been used successfully for parallel transcriptome and repertoire sequencing of single B and T cells, with dedicated bioinformatics pipelines named TraCeR^18^, BASIC^22^, BraCeR^19^, VDJPuzzle^21^. In FB5P-seq, the sequencing reads mainly cover the 5’-ends of mRNAs in a 3’ to 5’ orientation, which generates a broad coverage of the most 5’ half of mRNAs when using 67-nucleotide long reads. As a consequence, FB5P-seq can successfully reconstruct BCR or TCR repertoire sequences from the scRNA-seq reads when sequencing libraries on a 75-cycle Illumina flow cell at a targeted read depth of 5×10^5^ reads per cell, as done for the datasets presented here. In that configuration, the cost of performing FB5P-seq is roughly 5 € per cell (2.5 € library preparation + 2.5 € sequencing), which is significantly lower than Smart-Seq2 (30 € library preparation + 2.5-5 € sequencing)^11^. In FB5P-seq, only one library preparation is required per 96- well plate, which drastically reduces handling time when working with manual pipetting, as compared to generating one library per cell. Yet because library preparation only requires a small amount of pooled cDNA, the full-length amplified cDNA from each single-cell can be archived and used for downstream applications such as single-cell antibody cloning^34^.

Our bioinformatics pipeline for recovering repertoire sequences differs from previously published solutions^18,19,21,22^ in that it performs *de novo* transcriptome reconstruction on all sequences prior to filtering contigs corresponding to rearranged BCR or TCR chains. Because competing algorithms rely on full-length scRNA-seq paired-end reads, we did not test their ability to recover BCR or TCR sequences on FB5P-seq reads. Yet the demonstrated performance of our method on human B cell subsets, with close to 100% paired heavy and light chain reconstruction efficiency on PB/PCs, suggests it performs at least equally well as those methods.

We have designed the FB5P-seq TSOs with short 5-nucleotide UMIs, enabling the detection of a maximal molecular diversity of 4^5^=1,024 molecules for each gene per cell. Although this may be low for some genes in some cell types, in our analyses on lymphocytes, only the *CD74* gene showed recurrent saturating levels in Mem B cells and GC B cells (not shown). Longer UMI sequences could be used in the TSO design to prevent saturation issues^10,24^, but template switching efficiency may be affected by increased TSO length^35^. In our analyses of human lymphocytes, we have used very low levels of ERCC spike-ins (0.025 pg per well, corresponding to a 1:2,000,000 final dilution) compared to other scRNA-seq protocols^11,12,36^. We have optimized the concentration of ERCC spike-ins for having less than 5% of sequencing reads mapping on ERCC genes, to maximize the amount of biological information in our experiments while retaining the usefulness of ERCC spike-ins for quality control purposes. This would need to be adjusted when studying cell types with more mRNA content than primary lymphocytes.

The FB5P-seq protocol was designed to use FACS single-cell sorting for cell deposition into 96-well plates. Most cell sorting machines and software enable to use index sorting to record the flow cytometry parameters (cell size, fluorescence levels for each channel) of each sorted cell, including for markers that were not used in the gating strategy but were included during cell staining. This may be used to analyse mRNA / protein levels in parallel, or to investigate the relevance of cell surface markers to delineate transcriptionally robust cell subsets^7^. In our implementation of FB5P-seq, we use the 96-well plate format, sort single cells into 2μl lysis buffer and use manual pipetting throughout the protocol, but our method would only require minor adjustments to work on 384-well plates with smaller volumes and automatic liquid handlers like the MARS-seq protocol^37,38^. Furthermore, plates containing lysis buffer can be stored at −80°C and shipped on dry ice, before and after cell sorting, facilitating collaborative work and multisite projects.

High throughput droplet-based 5’-end scRNA-seq is an excellent option to analyse transcriptome and repertoire sequence for thousands of cells in parallel in complex tissues^23,39^. Combining 5’-end scRNA-seq with DNA-tagged antibody labelling and barcoding may further enable the multiplexing of several samples and the analysis of the surface phenotype of cells^40^. Yet, because those approaches best apply when many thousands of cells are available for input, we believe FB5P-seq is a valuable option to obtain the same multimodal information when focusing on rare cells defined by a complex surface phenotype. We expect our method will provide valuable insights for decoding the mechanisms regulating the molecular and functional diversity of lymphocytes during protective immunity, auto-immunity, cancer immunity, or lymphoma development and progression.

## Materials and Methods

### Human samples

Non-malignant tonsil samples from a 35-year old male (Tonsil 1) and a 30-year old female (Tonsil 2) were obtained as frozen live cell suspensions from the CeVi collection of the Institute Carnot/Calym (ANR, France, https://www.calym.org/-Viable-cell-collection-CeVi-.html).

Peripheral blood mononuclear cells (PBMCs) were collected in Nantes University Hospital and used fresh in peptide restimulation assays for isolating C.alb-specific T cells.

Written informed consent was obtained from the donors.

### Flow cytometry and cell sorting of B cell subsets

Frozen live cell suspensions were thawed at 37°C in RPMI + 10% FCS, then washed and resuspended in FACS buffer (PBS + 5% FCS + 2 mM EDTA) at a concentration of 10^8^ cells/ml for staining. Cells were first incubated with 2% normal mouse serum and Fc-Block (BD Biosciences) for 10 min on ice. Then cells were incubated with a mix of fluorophore-conjugated antibodies (**Table S2**) for 30 min on ice. Cells were washed in PBS, then incubated with the Live/Dead Fixable Aqua Dead Cell Stain (Thermofisher) for 10 min on ice. After a final wash in FACS buffer, cells were resuspended in FACS buffer at a concentration of 10^7^ cells/ml for cell sorting on a 4-laser BD FACS Influx (BD Biosciences).

Mem B cells were gated as CD3^-^CD14^--^IgD^-^CD20^+^CD10^-^CD38^lo^CD27^+^SSC^lo^ single live cells. GC B cells were gated as CD3^-^CD14^-^IgD^-^CD20^+^CD10^+^CD38^+^ single live cells. PB/PC cells were gated as CD3^-^CD14^-^IgD^-^CD38^hi^CD27^+^SSC^hi^ single live cells.

### Restimulation and cell sorting of antigen-specific T cells

Fresh PBMCs (10-20×10^6^ cells, final concentration 10×10^6^ cells/ml) were stimulated for 3h at 37°C with 0.6 nmol/ml PepTivator *Candida albicans* MP65 (pool of 15 amino acids length peptides with 11 amino acid overlap, Miltenyi Biotec) in RPMI + 5% human serum in the presence of 1 μg/ml anti-CD40 (HB14, Miltenyi Biotec). After stimulation, PBMCs were labeled with PE-conjugated anti-CD154 (5C8, Miltenyi Biotec) and enriched with anti-PE magnetic beads (Miltenyi Biotec)^41^. After enrichment, cells were stained with PerCP-Cy5.5 anti-CD4 (RPA-T4, Biolegend), AlexaFluor700 anti-CD3 (SK7, Biolegend) and APC-Cy7 anti-CD45RA (HI100, Biolegend), and antigen-specific T cells were gated as CD3^+^CD4^+^CD45RA^-^CD154^+^ single live cells for single-cell sorting.

### Single-cell RNA-seq

Single cells were FACS sorted into ice-cold 96-well PCR plates (Thermofisher) containing 2 μl lysis mix per well. The lysis mix contained 0.5 μl 0.4% (v/v) Triton X-100 (Sigma-Aldrich), 0.05 μl 40 U/μl RnaseOUT (Thermofisher), 0.08 μl 25 mM dNTP mix (Thermofisher), 0.5 μl 10 μM (dT)30_Smarter primer^8^ (**Table S3**), 0.05 μl 0.5 pg/μl External RNA Controls Consortium (ERCC) spike-ins mix (Thermofisher), and 0.82 μl PCR-grade H_2_0 (Qiagen).

For B cell subsets sorting, the index-sorting mode was activated to record the different fluorescence intensity of each sorted single-cell. Index-sorting FCS files were visualized in FlowJo software and compensated parameters values were exported in CSV tables for further processing. For visualization on linear scales in the R programming software, we applied the hyperbolic arcsine transformation on fluorescence parameters. In every 96-well plate, two wells (H1, H12) were left empty and processed throughout the protocol as negative controls.

Immediately after cell sorting, each plate was covered with adhesive film (Thermofisher), briefly spun down in a benchtop plate centrifuge, and frozen on dry ice. Plates containing single cells in lysis mix were stored at −80°C and shipped on dry ice (only T cells) until further processing.

The plate containing single cells in lysis mix was thawed on ice, briefly spun down in a benchtop plate centrifuge, and incubated in a thermal cycler for 3 minutes at 72°C (lid temperature 72°C). Immediately after, the plate was placed back on ice and 3 μl RT mastermix was added to each well. The RT mastermix contained 0.25 μl 200 U/μl SuperScript II (Thermofisher), 0.25 μl 40 U/μl RnaseOUT (Thermofisher), and 2.5 μl 2x RT mastermix. The 2x RT mastermix contained 1 μl 5x SuperScript II buffer (Thermofisher), 0.25 μl 100 mM DTT (Thermofisher), 1 μl 5 M betaine (Sigma-Aldrich), 0.03 μl 1 M MgCl_2_ (Sigma-Aldrich), 0.125 μl 100 μM well-specific template switching oligonucleotide TSO_BCx_UMI5_TATA (**Table S3** and **Table S4**), and 0.095 μl PCR-grade H_2_O (Qiagen). Reverse transcription was performed in a thermal cycler (lid temperature 70°C) by 90 min at 42°C, followed by 10 cycles of 2 min at 50°C and 2 min at 42°C, then 15 min at 70°C. Plates with single-cell cDNA were stored at −20°C until further processing.

For cDNA amplification, 7.5 μl LD-PCR mastermix were added to each well. The LD-PCR mastermix contained 6.25 μl 2x KAPA HiFi HotStart ReadyMix (Roche Diagnostics), 0.125 μl 20 μM PCR_Satija forward primer^9^ (**Table S3**), 0.125 μl 20 μM SmarterR reverse primer^8^ (**Table S3**), and 1 μl PCR-grade H_2_O (Qiagen). The amplification was performed in a thermal cycler (lid temperature 98°C) by 3 min at 98°C, followed by 22 cycles of 15 sec at 98°C, 20 sec at 67°C, 6 min at 72°C, then a final elongation for 5 min at 72°C. Plates with amplified single-cell cDNA were stored at -20°C until further processing.

For library preparation, 5 μl amplified cDNA from each well of a 96-well plate were pooled and completed to 500 μl with PCR-grade H_2_O (Qiagen). Two rounds of 0.6X solid-phase reversible immobilization beads (AmpureXP, Beckman, or CleanNGS, Proteigene) cleaning were used to purify 100 μl pooled cDNA with final elution in 15 μl PCR-grade H_2_O (Qiagen). After quantification with Qubit dsDNA HS assay (Thermofisher), 800 pg purified cDNA pool were processed with the Nextera XT DNA sample Preparation kit (Illumina), according to the manufacturer’s instructions with modifications to enrich 5’-ends of tagmented cDNA during library PCR. After tagmentation and neutralization, 25 μl tagmented cDNA was amplified with 15 μl Nextera PCR Mastermix, 5 μl Nextera i5 primer (S5xx, Illumina), and 5 μl of a custom i7 primer mix^9^ (0.5 μM i7_BCx + 10 μM i7_primer, **Table S3**). The amplification was performed in a thermal cycler (lid temperature 72°C) by 3 min at 72°C, 30 sec at 95°C, followed by 12 cycles of 10 sec at 95°C, 30 sec at 55°C, 30 sec at 72°C, then a final elongation for 5 min at 72°C. The resulting library was purified with 0.8X solid-phase reversible immobilization beads (AmpureXP, Beckman, or CleanNGS, Proteigene).

Libraries generated from multiple 96-well plates of single cells and carrying distinct i7 barcodes were pooled for sequencing on an Illumina NextSeq550 platform, with High Output 75 cycles flow cells, targeting 5x10^5^ reads per cell in paired-end single-index mode with the following primers (**Table S3**) and cycles: Read1 (Read1_SP, 67 cycles), Read i7 (i7_SP, 8 cycles), Read2 (Read2_SP, 16 cycles).

### Single-cell RNA-seq data processing

We used a custom bioinformatics pipeline to process fastq files and generate single-cell gene expression matrices and BCR or TCR sequence files. Detailed instructions for running the FB5P-seq bioinformatics pipeline can be found at https://github.com/MilpiedLab/FB5P-seq. Briefly, the pipeline to obtain gene expression matrices was adapted from the Drop-seq pipeline^26^, relied on extracting the cell barcode and UMI from Read2 and aligning Read1 on the reference genome using STAR and HTSeqCount. For BCR or TCR sequence reconstruction, we used Trinity for *de novo* transcriptome assembly for each cell based on Read1 sequences, then filtered the resulting isoforms for productive BCR or TCR sequences using MigMap, Blastn and Kallisto. Briefly, MigMap was used to assess whether reconstructed contigs corresponded to a productive V(D)J rearrangement and to identify germline V, D and J genes and CDR3 sequence for each contig. For each cell, reconstructed contigs corresponding to the same V(D)J rearrangement were merged, keeping the largest sequence for further analysis. We used Blastn to align the reconstructed BCR or TCR contigs against reference sequences of constant region genes, and discarded contigs with no constant region identified in-frame with the V(D)J rearrangement. Finally, we used the pseudoaligner Kallisto to map each cell’s FB5P-seq Read1 sequences on its reconstructed contigs and quantify contig expression. In cases where several contigs corresponding to the same BCR or TCR chain had passed the above filters, we retained the contig with the highest expression level.

The per well accuracy (**Figure 3B**) was computed as the Pearson correlation coefficient between log_10_(UMI_ERCC-xxxxx_+1) and log_10_(#mol_ERCC-xxxxx_+1), where UMI_ERCC-xxxxx_ is the UMI count for gene *ERCC-xxxxx* in the well, and #mol_ERCC-xxxxx_ is the actual number of molecules for *ERCC-xxxxx* in the well (based on a 1:2,000,000 dilution in 2 μl lysis mix per well). For each well, only *ERCC-xxxxx* which were detected (UMI_ERCC-xxxxx_>0) were considered for calculating the accuracy.

To estimate sensitivity (**Figure 3C**), the percentage of wells with at least one molecule detected (UMI_ERCC-xxxxx_>0) was calculated over all the wells from 5 or 6 96-well plates corresponding to human B cells sorted from Tonsil 1 or Tonsil 2, respectively. The value for each *ERCC-xxxxx* gene was plotted against log_10_(#mol_ERCC-xxxxx_) and a standard curve was interpolated with asymmetric sigmoidal 5PL model in GraphPad Prism 8.1.2 to compute the EC50 for each dataset.

The normalized coverage over genes (**Figure 3F**) was computed with RSeQC *geneBody_coverage* on *bam* files from 11 scRNA-seq 96-well plates corresponding to human B cells sorted from Tonsil 1 and Tonsil 2.

### Single-cell gene expression analysis

Quality control was performed on each dataset (Tonsil 1, Tonsil 2, T cells) independently to remove poor quality cells. Cells with less than 250 genes detected were removed. We further excluded cells with values below 3 median absolute deviations (MADs) from the median for UMI counts, for the number of genes detected, or for ERCC accuracy, and cells with values above 3 MADs from the median for ERCC transcript percentage.

For each cell, gene expression UMI count values were log-normalized with Seurat v3^42^ *NormalizeData* with a scale factor of 10,000. Data from B cells of Tonsil 1 and Tonsil 2 were analyzed together. Data from *C.alb*-specific T cells were analyzed separately. Four thousand variable genes, excluding BCR or TCR genes, were identified with Seurat *FindVariableFeatures*. After centering and scaling with Seurat *ScaleData*, principal component analysis was performed on variable genes with Seurat *RunPCA*, and embedded in two-dimensional t-SNE plots with Seurat *RunT-SNE* on 40 principal components. Cell cycle phases were attributed with Seurat *CellCycleScoring* using lists of S phase or G2/M phase specific genes as described^43^. Plots showing t-SNE embeddings colored by index sorting protein expression or other metadata (including BCR or TCR sequence related information) were generated with ggplot2 *ggplot*. Plots showing t-SNE embeddings colored by gene expression were generated by Seurat *FeaturePlot*. Gene expression heatmaps were plotted with a custom function (available upon request).

## Accession codes

The single-cell RNA-seq data generated in the current study are available in the Gene Expression Omnibus database under accession code GSE137275.

## Acknowledgements

We are grateful to P. Barbry, K. Lebrigand and M.-J. Arguel from Institut de Pharmacologie Moléculaire et Cellulaire for initial discussions on method design and bioinformatics analysis. We thank all members of the B. Nadel and P. Milpied laboratories at Centre d’Immunologie de Marseille-Luminy for useful discussions. We are grateful to collaborators who trusted us with our method before publication. We thank the Bioinformatics Core Facility of Centre d’Immunologie de Marseille-Luminy for helpful discussions and comments. We are thankful to E. Mollaret from Institut Carnot CALYM for organizing tonsil samples transfer from the CeVi biobank. We acknowledge HalioDX and the UCA Genomix platform for sequencing. This work was supported by grants from Fondation ARC, Cancéropôle Provence-Alpes-Côte d’Azur and ANR (JCJC MoDEx-GC) to P.M. This work was supported by institutional grants from INSERM, CNRS and Aix-Marseille University to the CIML. This work was granted access to the HPC resources of Aix-Marseille Université financed by the project Equip@Meso (ANR-10-EQPX-29-01) of the program « Investissements d’Avenir » supervised by the Agence Nationale de la Recherche.

## Author contributions

N.A. developed FB5P-seq, designed experiments, performed experiments, analyzed the data and wrote the manuscript. I.C.-M. developed FB5P-seq, designed the bioinformatics pipeline and analyzed the data. C.D. assembled and automatized the bioinformatics pipeline and analyzed the data. L.G. performed experiments. A.R. performed the human T cell experiment. L.S. supervised the development and assembly of the bioinformatics pipeline. P.M. developed FB5P-seq, supervised the study, designed experiments, analyzed the data and wrote the manuscript. All authors reviewed and approved the manuscript.

## Supplementary Figures

**Figure S1.**
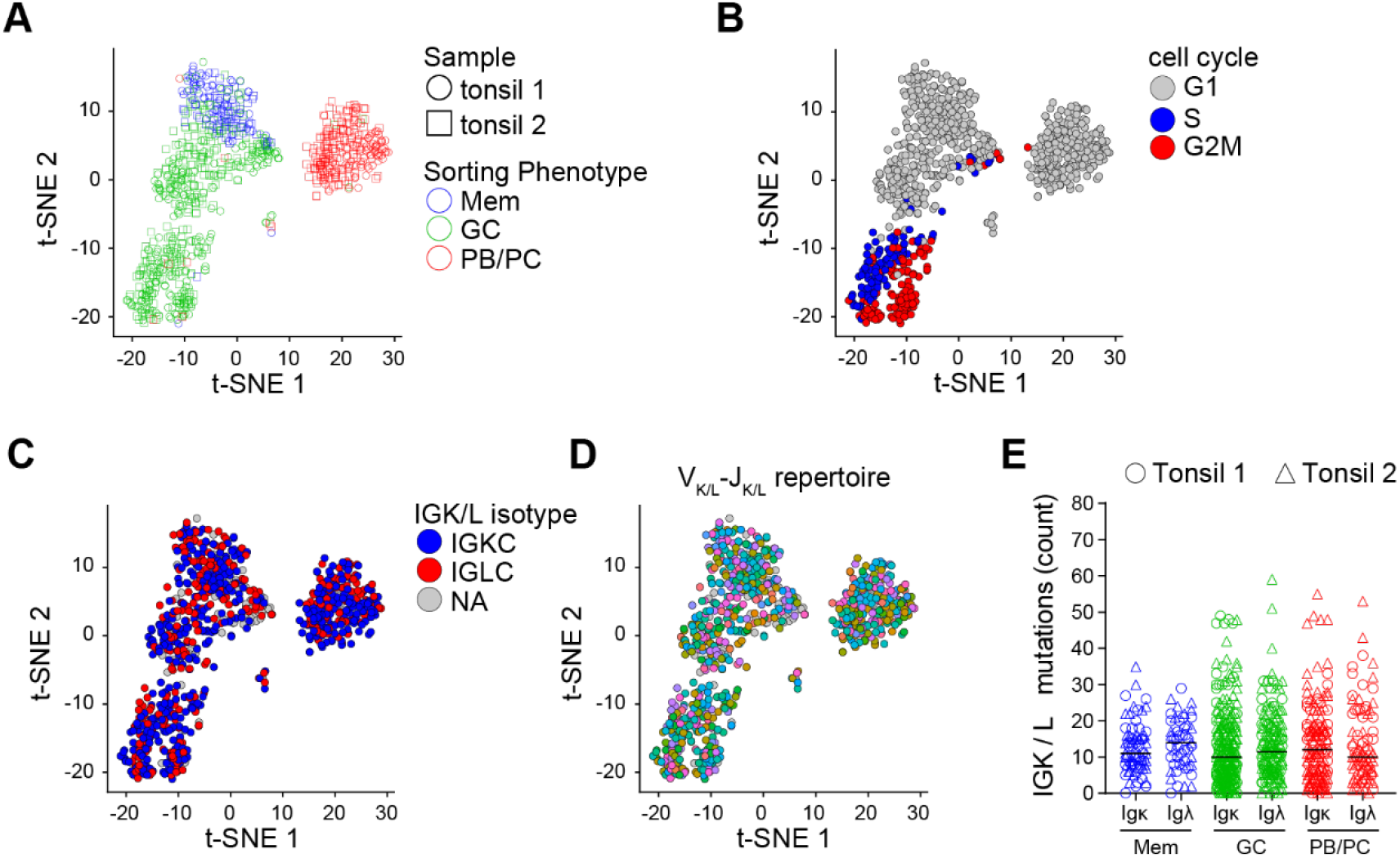
FB5P-seq analysis of human tonsil B cell subsets (related to Figure 4). **(A)** t-SNE map of single human B cell subsets from Tonsil 1 (circles) and Tonsil 2 (squares) computed on 4,000 variable genes excluding BCR genes. Cells are colored based on sorting phenotype (n=845 cells). **(B-D)** t-SNE map of single human tonsil B cells colored by cell cycle phase **(B)**, BCR light chain isotype (NA: not applicable i.e. no IGK/L reconstructed) **(C),** or V_K/L_-J_K/L_ repertoire (grey cells: no IGK/L reconstructed) **(D)**. **(E)** Scatter plots showing IGK/L mutation frequency in human Tonsil 1 (circles) and Tonsil 2 (triangles) B cells sorted by their IGK/L isotype and phenotype (Mem B cells: n=71 Igκ+, n=51 Igλ+; GC B cells: n=253 Igκ+, n=163 Igλ+; PB/PCS: n=139 Igκ+, n=84 Igλ+).

## Supplementary Tables

**Table S1.** Quality control characteristics of FB5P-seq datasets.

| **Sample** | **# plates** | **# cells sorted** | **# reads per cell (mean)** | **# cells after QC** | **% cells passing QC** |
| --- | --- | --- | --- | --- | --- |
| Tonsil 1 B cells | 5 | 420 | 540,087 | 386 | 91.9 |
| Tonsil 2 B cells | 6 | 504 | 525,890 | 459 | 91.1 |
| CD4 T cells | 1 | 84 | 397,670 | 82 | 97.6 |

**Table S2.**
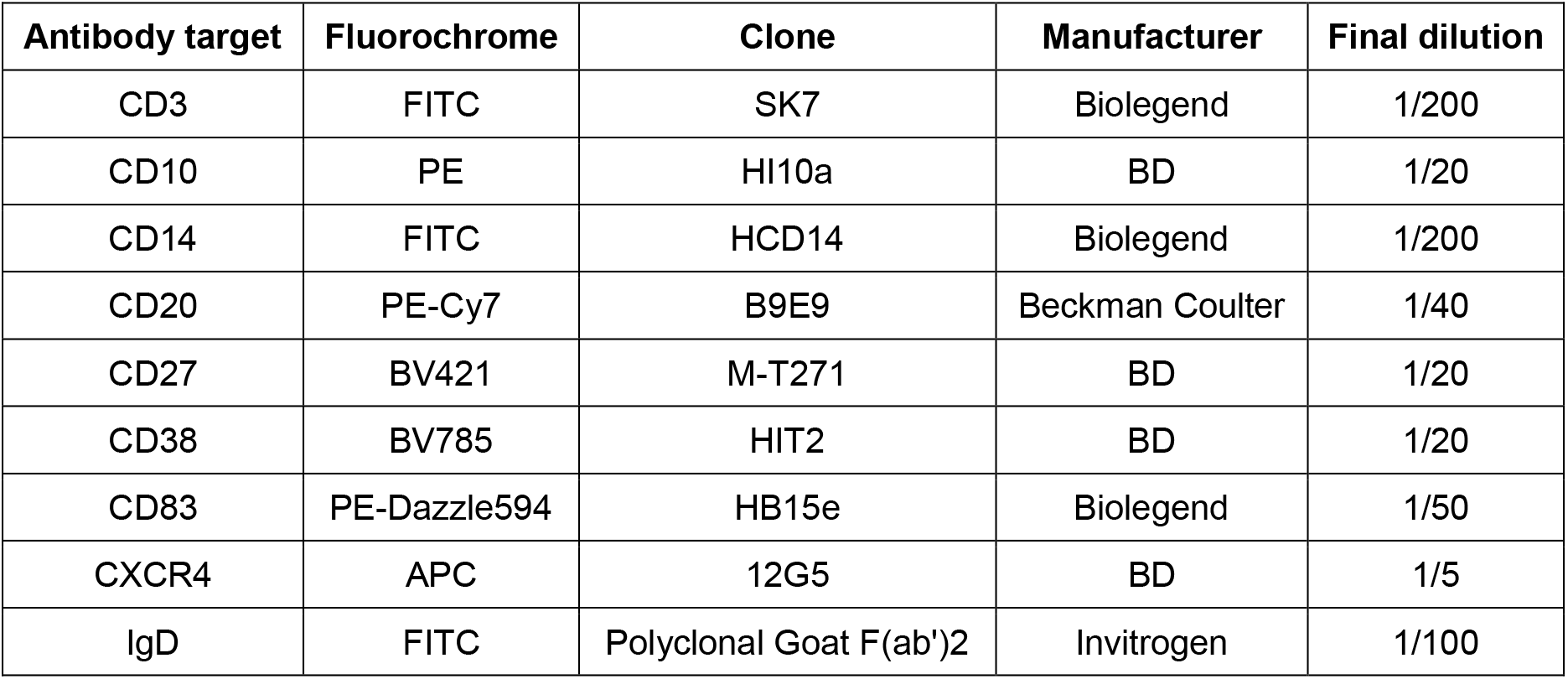
List of antibodies for FACS analysis of tonsil B cells.

**Table S3.**
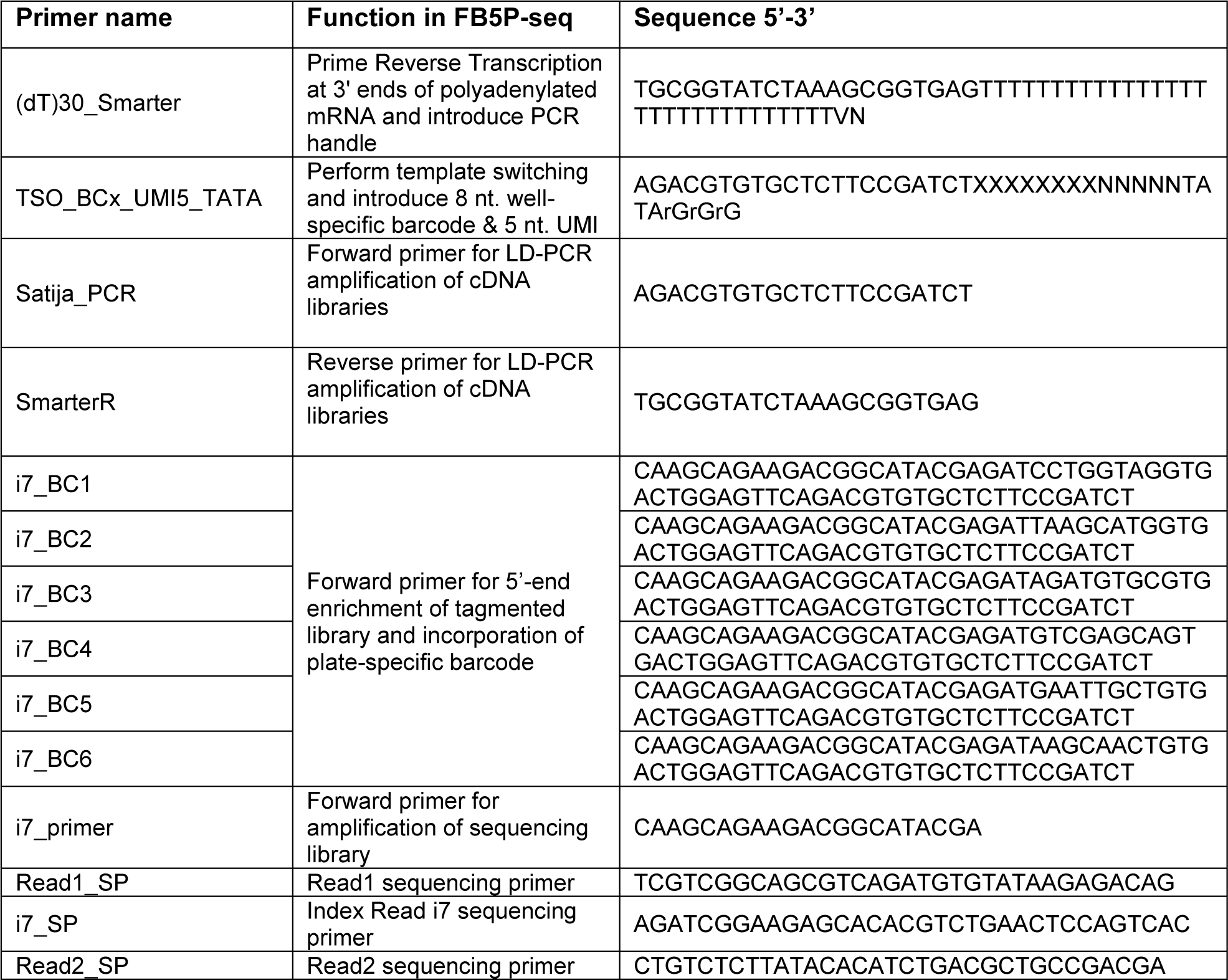
Primers used in FB5P-seq.

**Table S4.**
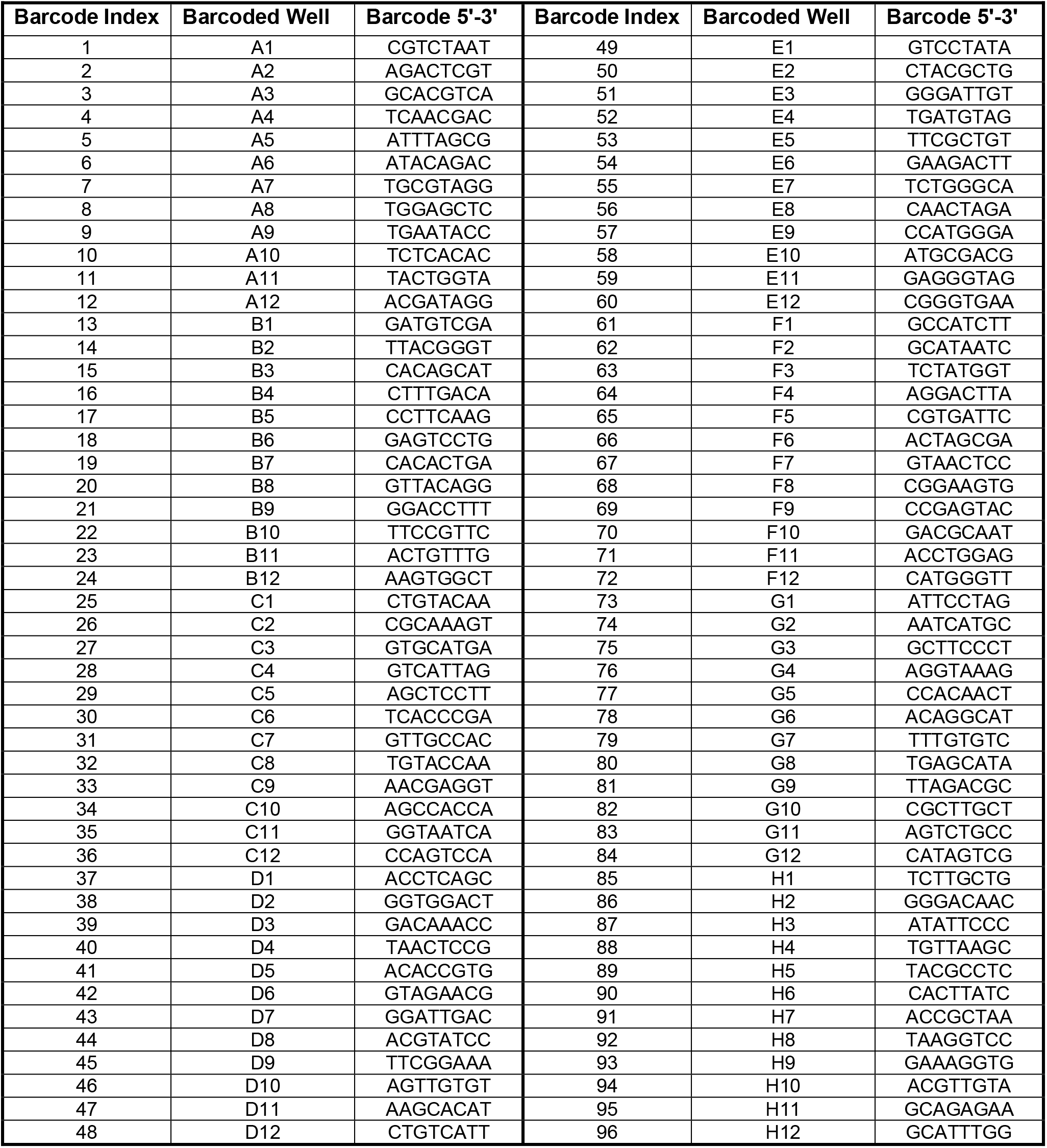
List of well-specific barcodes used in TSO_BCx_UMI5_TATA primers.

